# The PIP4K2 inhibitor THZ-P1-2 exhibits antileukemia activity by disruption of mitochondrial homeostasis and autophagy

**DOI:** 10.1101/2022.08.20.504641

**Authors:** Keli Lima, Diego Antonio Pereira-Martins, Lívia Bassani Lins de Miranda, Juan Luiz Coelho-Silva, Giovana da Silva Leandro, Isabel Weinhäuser, Rita de Cássia Cavaglieri, Aline Medeiros de Leal, Wellington Fernandes da Silva, Ana Paula Alencar de Lima Lange, Elvira Deolinda Rodrigues Pereira Velloso, Emmanuel Griessinger, Jacobien R Hilberink, Gerwin Huls, Jan Jacob Schuringa, Eduardo Magalhães Rego, João Agostinho Machado-Neto

**Affiliations:** Laboratory of Medical Investigation in Pathogenesis and Targeted Therapy in Onco-Immuno-Hematology (LIM-31), Department of Internal Medicine, Hematology Division, Faculdade de Medicina, University of São Paulo, São Paulo, Brazil; Department of Pharmacology, Institute of Biomedical Sciences, University of São Paulo, São Paulo, Brazil; Department of Medical Imaging, Hematology, and Oncology, Ribeirão Preto Medical School, University of São Paulo, Ribeirão Preto, Brazil; Department of Experimental Hematology, Cancer Research Center Groningen, University Medical Center Groningen, University of Groningen, Groningen, Netherlands; Department of Microbiology, Institute of Biomedical Sciences, University of São Paulo, São Paulo, Brazil

**Keywords:** Acute leukemia, Phosphatidylinositol-5-phosphate 4-kinase type 2, THZ-P1-2, Mitochondrial homeostasis, Autophagy

## Abstract

Treatment of acute leukemia is challenging due to genetic heterogeneity between and even within patients. Leukemic stem cells (LSCs) are relatively drug-resistant and frequently lead to relapse. Their plasticity and capacity to adapt to extracellular stress, in which mitochondrial metabolism and autophagy play important roles, further complicates treatment. Genetic models of phosphatidylinositol-5-phosphate 4-kinase type 2 proteins (PIP4K2s) inhibition demonstrated the relevance of these enzymes in mitochondrial homeostasis and autophagic flux. Here, we uncover the cellular and molecular effects of THZ-P1-2, a pan-inhibitor of PIP4K2s, in acute leukemia cells. THZ-P1-2 reduced cell viability and induced DNA damage, apoptosis, loss of mitochondrial membrane potential, and accumulation of acidic vesicular organelles. Protein expression analysis revealed that THZ-P1-2 impaired autophagy flux. In addition, THZ-P1-2 induced cell differentiation and showed synergistic effects with venetoclax in resistant leukemic models. In primary leukemia cells, LC-MS/MS-based proteome analysis revealed that sensitivity to THZ-P1-2 was associated with mitochondrial metabolism, cell cycle, cell-of-origin, and the TP53 pathway. Minimal effects of THZ-P1-2 observed in healthy CD34^+^ cells suggested a favorable therapeutic window. Our study provides insight into pharmacological inhibition of PIP4Ks targeting mitochondrial homeostasis and autophagy shedding light on a new class of drugs for acute leukemias.

## Introduction

Acute leukemias comprise a heterogeneous group of highly aggressive hematological malignancies. In adults, both acute myeloid leukemia (AML) and acute lymphoblastic leukemia (ALL) have high rates of relapse and poor prognosis [1,2]. Significant advances have been made in understanding the molecular basis of these diseases, which has allowed the identification of diagnostic markers as well as prognostic factors and novel FDA-approved targeted therapies [3,4,5]. Nevertheless, the therapeutic repertoire remains limited. Recent studies have indicated a prominent role for mitochondrial metabolism in the therapeutic response of these neoplasms, revealing increased sensitivity of leukemic stem cells (LSC) to disruptions of oxidative phosphorylation (OXPHOS) process [6,7]. These vulnerabilities may be specifically targeted with the BCL2 inhibitor venetoclax, which suppresses the tricarboxylic acid (TCA) cycle by inhibiting amino acid metabolism [8,9,10]. Additionally, autophagy and lysosomal biosynthesis have emerged as potential targets to improve the therapeutic response of these diseases [11,12].

The family of phosphatidylinositol-5-phosphate 4-kinase type 2 proteins (PIP4K2s) comprises three lipid kinase members (PIP4K2A, PIP4K2B, and PIP4K2C) that phosphorylate the phosphatidylinositol (PI)5P at position four of the inositol ring, generating PI-4,5-P_2_ [13]. Furthermore, these enzymes have been associated with the local and specialized generation of PI-4,5-P2 pools in intracellular membranes, being critical for the functionality of endosomes, autophagosomes, lysosomes, and peroxisomes [14,15,16,17].

Under physiological conditions, PI-4,5-P_2_-mediated signaling is implicated in many cellular processes, including proliferation, survival, glucose uptake, and cytoskeletal organization, which have been reported to be deregulated in breast cancer, acute leukemias, glioblastoma, and soft tissue sarcomas [13,18]. Cancer genomic and transcriptomic landscape studies reveal that PIP4K2-related genes are usually not mutated, but that their expression is frequently altered [13]. In ALL, single nucleotide polymorphisms (SNPs) in regulatory regions of the *PIP4K2A* gene were associated with increased mRNA levels and risk of disease development [19,20]. In hematological malignancies, PIP4K2A was identified as an essential protein for proliferation, clonogenicity, and survival of leukemia-initiating cells [21]. In AML patients, high *PIP4K2A* and *PIP4K2C* transcript levels were associated with adverse cytogenetic risk and poor clinical outcome [22,23].

THZ-P1-2 is the prototype of a new class of drugs that acts as a selective pan-inhibitor of PIP4K2s. This drug covalently binds to a single cysteine residue located in a disordered loop close to the ATP binding site of the kinase domain present in the three PIP4K2 isoforms, which irreversibly inhibits the activity of these enzymes. Genetic inhibition of PIP4K2s indicated that inhibition of at least two isoforms (PIP4K2A/B) is necessary to obtain a pronounced biological effect [16,24,25], suggesting that selective pharmacological inhibitors for more than one isoform are of interest. Initial studies showed that THZ-P1-2 reduces cell viability by inhibiting autophagic flux in solid tumors [26], but efficacy in acute leukemias has not yet been explored. Here, we show that THZ-P1-2 provides potent anti-leukemic effects in AML and ALL, and we uncover the underlying molecular mechanisms.

## Material and methods

### Patients

Bone marrow or peripheral blood samples from AML and ALL patients were used after informed consent and protocol approved by the Ethical Committee of the Institute of Biomedical Sciences of the University of São Paulo (CAAE: 39510920.1.0000.5467) or the medical ethical committee of the UMCG in accordance with the Declaration of Helsinki. Mononuclear cells were isolated by density gradient separation (Ficoll, Sigma-Aldrich, Sant Louis, MO, USA) according to the manufactory’s instructions. An overview of patients’ characteristics is described in Supplementary Table 1-3. Neonatal cord blood (CB) samples were obtained from healthy full-term pregnancies in accordance with the Declaration of Helsinki at the obstetrics departments at the Martini Hospital and University Medical Center Groningen. CD34^+^ cells were isolated using an autoMACS hematopoietic progenitor magnetic-associated cell-sorting kit (#130-046-703, Miltenyi Biotech, Bergisch Gladbach, Germany) according to the manufacturer’s instructions.

### Cell lines and inhibitors

OCI-AML3, Kasumi-1, MOLM-13, and MV4-11 cells were obtained from Deutsche Sammlung von Mikroorganismen und Zellkulturen (DSMZ, Germany). THP-1 and HL-60 cells were obtained from American Type Culture Collection (ATCC, USA). NB4 cells were kindly provided by Prof. Pier Paolo Pandolfi (Beth Israel Deaconess Medical Center, Havard Medical School, Boston, USA). U-937, K-562, KU812, HEL, Jurkat, Namalwa, Daudi, and Raji cells were kindly provided by Prof. Sara Teresinha Olalla Saad (University of Campinas, Campinas, Brazil). CEM, NALM6, and REH were kindly provided by Gilberto Carlos Franchi Junior (University of Campinas, Campinas, Brazil). SET-2 cells were kindly provided by Prof. Fabíola Attié de Castro (University of São Paulo, Ribeirão Preto, Brazil). SUP-B15 was kindly provided by Dr. Lucas Eduardo Botelho de Souza (National Institute of Science and Technology in Stem Cells and Cell Therapy, Ribeirão Preto, Brazil). MOLT-4 cells were kindly provided by Prof. Gisele Monteiro (University of São Paulo, São Paulo, Brazil). Cells were cultured in appropriate media (RPMI-1640, IMDM, or alpha-MEM) supplemented with 10 or 20% fetal bovine serum (FBS) according to the ATCC and DSMZ recommendations, plus 1% penicillin/streptomycin and maintained at 5% CO_2_ and 37°C. THZ-P1-2 was obtained from MedChemExpress (Monmouth Junction, NJ, USA) and diluted to a 25 mM stock solution in dimethyl sulfoxide (Me_2_SO_4_; DMSO). Venetoclax (ABT-199) was obtained from TargetMol (Target Molecule Corp., Boston, MA, USA) and diluted to a 50 mM stock solution in DMSO. Cytarabine (AraC) was obtained from Accord (Accord Healthcare B.V., Utrecht, Netherlands) and diluted to a 1 mM stock solution in DMSO.

### Cell viability assay

In total 2 × 10^4^ cells per well were seeded in a 96-well plate in the appropriate medium in the presence of vehicle or different concentrations of THZ-P1-2 (1.6, 3.2, 6.4, 12.5, 25, 50, or 100 µM) for 24, 48, and/or 72 h. Next, 10 μl methylthiazoletetrazolium (MTT, Sigma-Aldrich) solution (5 mg/mL) was added and incubated at 37 °C, 5% CO_2_ for 4 h. The reaction was stopped using 100 μL 0.1N HCl in anhydrous isopropanol. Cell viability was evaluated by measuring the absorbance at 570 nm. IC_50_ values were calculated using nonlinear regression analysis in GraphPad Prism 5 (GraphPad Software, Inc., San Diego, CA, USA).

### Cell cycle assay

In total 6 × 10^5^ cells per well were seeded in 6-well plates supplemented with vehicle or THZ-P1-2 (1.6, 3.2, and 6.4 μM), harvested at 24 h, fixed with 70% ethanol, and stored at 4°C for at least 2 h. Next, the fixed cells were stained with 20 μg/mL propidium iodide (PI) containing 10 μg/mL RNase A for 30 min at room temperature in a light-protected area. DNA content distribution was acquired using flow cytometry (FACSCalibur; Becton Dickinson, Franklin Lakes, NJ, USA) and analyzed using FlowJo software (Treestar, Inc., San Carlos, CA, USA).

### Comet assay

For the alkaline comet assay, MV4-11 and Jurkat cells (1.5 × 10^5^ per 35 mm cell culture dish) were treated with vehicle or THZ-P1-2 (3.2 and 6.4 µM) for 48 h. Next, cells were counted and suspended at a concentration of 1 × 10^5^ cells per mL. A total of 12 µL of cell solution was suspended in low melting point (0.5%) agarose suspension in PBS at 37°C and solutions were added onto pre-coated CometAssay® Kit slides (Trevigen, Gaithersburg, MD, USA). After solidifying of agarose, slides were immersed in a cold lysis solution (CometAssay Lysis Solution, Trevigen) for at least 40 min, protected from light. Still, under the protection of light, the slides were immersed in an alkaline solution (300 mM NaOH and 1 mM EDTA, pH 13.0) for 20 minutes to allow the unfolding of the DNA structure. Then, the slides were arranged horizontally in the electrophoresis vat with an alkaline solution (30 mM NaOH and 1 mM EDTA, pH 13.0) at 4°C. Electrophoresis was delivered at 300 mA at 4°C for 30 min. After the electro, the slides were again neutralized with dH2O for 5 min and fixed with 70% ethanol. For DNA staining, SYBR™ Green I Nucleic Acid Gel Stain (Invitrogen Molecular Probes, Eugene, OR, USA) was used for 5 min. Slides were analyzed under a Zeiss fluorescence microscope (Axiovert 200, Zeiss, Jena, Germany) with a 510–560 nm filter and a 590 nm barrier. The DNA head-to-tail ratio was scored from 100 comets on each slide analyzed using the LUCIA Comet Assay™ software (Laboratory Image, Prague, Czech Republic).

### Apoptosis assay

In total 1 × 10^5^ cells per well were seeded in a 24-well plate in FBS-supplemented appropriate media in the presence of vehicle or THZ-P1-2 (1.6, 3.2, and 6.4 μM) for 24 h. Next, the cells were washed with ice-cold phosphate-buffered saline (PBS) and resuspended in a binding buffer containing 1 μg/mL propidium iodide (PI) and 1 μg/mL APC-labeled annexin V (BD Pharmingen, San Diego, CA, USA). All specimens were analyzed by flow cytometry (FACSCalibur) after incubation for 15 min at room temperature in a light-protected area. Ten thousand events were acquired for each sample.

### Mitochondrial membrane potential evaluation

In total 1 × 10^5^ cells per well were seeded in a 24-well plate in FBS-supplemented appropriate media in the presence of vehicle or THZ-P1-2 (1.6, 3.2, and/or 6.4 μM) for 24 h. Cells were then washed with PBS and resuspended in buffer assay red containing 5 μg/mL JC-1 (BD) and 10 000 events were acquired by flow cytometry (FACSCalibur) after incubation for 15 minutes at 37°C and 5% CO_2_ in a light-protected area. Alternatively, cells were stained with 100 nM tetramethylrhodamine, ethyl ester, perchlorate (TMRE) according to the manufactory’s instructions (Thermo Fisher Scientific, Waltham, MA, USA), and 10 000 events were acquired by flow cytometry using the BD LSR™II (BD Biosciences). The events were analyzed using FlowJo software (Treestar, Inc.).

### Acidic vesicular organelles staining by acridine orange

In total 1 × 10^5^ cells per well were seeded in a 24-well plate in FBS-supplemented appropriate media in the presence of vehicle or THZ-P1-2 (1.6, 3.2, and 6.4 μM) for 24 h. Cells were then washed with PBS and resuspended in PBS containing 0.1 μg/mL acridine orange (Sigma-Aldrich) and 10 000 events were acquired by flow cytometry (FACSCalibur) after incubation for 30 minutes at room temperature in a light-protected area and one wash with PBS. The events were analyzed using FlowJo software (Treestar, Inc.). The recommendations published by Thomé *et al*. [27] were followed during the acquisition and analysis of the data to avoid false-positive results. Alternately, OCI-AML3 and NALM6 cells stained with PBS containing 0.1 μg/mL acridine orange were evaluated for GFP and RFP channels using a fluorescent microscope (Lionheart FX Automated microscope; Agilent BioTek Instruments Inc., Santa Clara, CA, USA; magnification, ×400).

### Western blotting

Cells were treated with vehicle or THZ-P1-2 (1.6, 3.2, or 6.4 μM) for 24 h and submitted to total protein extraction using a buffer containing 100 mM Tris (pH 7.6), 1% Triton X-100, 2 mM PMSF, 10 mM Na_3_VO_4_, 100 mM NaF, 10 mM Na_4_P_2_O_7_, and 4 mM EDTA. Equal amounts of protein (30 μg) of the samples were subsequently subjected to SDS-PAGE in an electrophoresis device, followed by electrotransfer of the proteins to nitrocellulose membranes. The membranes were blocked with 5% non-fat dry milk and then incubated with specific primary antibodies diluted in blocking buffer and next with secondary antibodies conjugated to HRP (horseradish peroxidase). Western blot analysis was performed using a SuperSignal™ West Dura Extended Duration substrate system (Thermo Fisher Scientific) and a G: BOX Chemi XX6 gel document system (Syngene, Cambridge, UK). Antibodies directed against PARP1 (#9542), γH2AX (#9718), SQSTM1/p62 (#88588), LC3B (#2775), and α-tubulin (#2144) were obtained from Cell Signaling Technology (Danvers, MA, USA).

### Quantitative RT-PCR (qRT-PCR)

Cells were treated with vehicle or THZ-P1-2 (6.4 μM) for 24 h, after which total RNA was extracted using TRIzol reagent (Thermo Fisher Scientific). cDNA was synthesized from 1 μg RNA using a High-Capacity cDNA Reverse Transcription Kit (Thermo Fisher Scientific). Quantitative PCR (qPCR) was performed using a QuantStudio 3 Real-Time PCR System in conjunction with a SybrGreen System (ThermoFisher Scientific) for assessment of the expression of cell cycle-, apoptosis-, and autophagy-related genes (Supplementary Table 4). *HPRT1* and *ACTB* were used as reference genes. Relative quantification values were calculated using the 2^-ΔΔCT^ equation [28]. A negative ‘No Template Control’ was included for each primer pair. Data were illustrated using Morpheus (https://software.broadinstitute.org/morpheus/).

### Ex vivo assays

For cell viability evaluation by MTT assay, 2 × 10^5^ mononuclear primary cells per well were seeded in a 96-well plate in appropriate medium (AML samples: 30% FBS RPMI, 10 ng/mL IL3, 10 ng/mL FLT3L, 100 ng/mL TPO, 100 ng/mL SCF; ALL samples: 30% FBS RPMI, 30 ng/mL IL3, 100 ng/mL IL7, 100 ng/mL FLT3L, and 30 ng/mL SCF) in the presence of vehicle or different concentrations of THZ-P1-2 (1.6, 3.2, 6.4, 12.5, 25, 50, or 100 µM) for 72 h and submitted to analysis as described above. All cytokines and growth factors were acquired from PeproTech (Rocky Hill, NJ, USA). IC50 values were calculated using nonlinear regression analysis in GraphPad Prism 5.

Apoptosis and mitochondrial membrane potential analysis were performed using FITC-labeled annexin V/DAPI and TMRE staining, respectively, and flow cytometry analysis in gated human Sca1^-^ (to exclude the MS5 contamination), CD45^+^ (APCCy7, BD biosciences), CD34^+^ (or CD117^+^ for *NPM1*-mutant AMLs) cells. Briefly, mouse bone marrow-derived MS5-stromal cells were initially plated on gelatin-coated culture 24-well plates and expanded to form a confluent layer, to which primary AML mononuclear cells were added. In co-culture assays, primary AML cells were cultured in 12.5% FBS and 12.5% horse serum IMDM (Thermo Fisher Scientific), 1% penicillin and streptomycin (Life Technologies, Grand Island, USA), and 57.2 µM β-mercaptoethanol (Merck Sharp & Dohme BV, Haarlem, Netherlands), with the addition of 20 ng/mL G-CSF (Amgen), N-Plate (clinical grade TPO) (Amgen) and IL-3 (Sandoz). Cells were exposure to vehicle, AraC (250 and 500 nM), venetoclax (100 and 500 nM) and/or THZ-P1-2 (3.2 and 6.4 µM) for 72 hours. AraC and venetoclax were used as control (AML-related cytotoxic drugs).

For growth curves, CB CD34^+^ cells were expanded on the co-culture system with MS5-stromal cells in the presence of vehicle and THZ-P1-2 (3.2 and 6.4 µM). For colony formation assays, a total of 300 CB CD34^+^ cells were seeded on methylcellulose (H4230, Stem Cell Technologies, Vancouver, Canada) supplemented with SCF, FLT3L, N-plate (all 100 ng/mL), and EPO, IL-3 and IL-6 (all 20 ng/mL) in the presence of vehicle or THZ-P1-2 (3.2 and 6.4 µM). After 8 days for CFU-E/BFU-E and 14 days for CFU-G/GM, colonies were identified and counted. All cell cultures were grown at 37°C and 5% CO_2_.

### Proteome and gene set enrichment analysis (GSEA)

Proteome studies were performed as previously described (available at PRIDE under PXD030463) [29]. Gene Set Enrichment Analysis (GSEA) was performed with GSEA v.4.0 [30] using the following MSigDB curated genesets: Hallmarks, Reactome, and Kegg. Enrichment scores (ES) were calculated based on the Kolmogorov-Smirnov statistic, tested for significance using 1000 permutations, and normalized (NES) to take into account the size of each gene set. A significance cutoff of FDR q-values < 0.25 was used.

### Oxygen consumption and extracellular acidification rate measurements

Oxygen consumption rate (OCR) and extracellular acidification rate (ECAR) were measured using the Seahorse XF96 analyzer (Seahorse Bioscience, Agilent, US) at 37 °C. A total of 1 × 10^5^ cells (treated with vehicle or 6.4 µM THZ-P1-2 for 24h) were seeded per well in poly-L-lysine (Sigma-Aldrich) coated Seahorse XF96 plates in 200 μL XF Assay Medium (Modified DMEM, Seahorse Bioscience). For OCR measurements, XF Assay Medium was supplemented with 10 mM glucose. Oligomycin A (2.5 µM), FCCP (carbonyl cyanide-4-(trifluorometh oxy) phenylhydrazone) (2.5 µM), and 2 µM antimycin (2 µM) plus rotenone (2 µM) were sequentially injected in 25 µL volume to measure basal and maximal OCR levels (all reagents from Sigma-Aldrich). For ECAR measurements, a Glucose-free XF Assay medium was added to the cells and 10 mM Glucose, 2.5 µM oligomycin A, and 100 mM 2-deoxy-D-glucose were sequentially injected in 25 µL volume (all reagents from Sigma-Aldrich). All XF96 protocols consisted of 4 times mix (2 min) and measurement (2 min) cycles, allowing for determination of OCR at basal and also in between injections. Both basal and maximal OCR levels were calculated by assessing the metabolic response of the cells following the manufacturer’s suggestions. The OCR measurements were normalized to the viable number of cells used for the assay.

### Cell differentiation analysis

OCI-AML3, THP-1, and NB4 cells were exposed to vehicle or THZ-P1-2 (1.6, 3.2, or 6.4 μM) for 72 h. Next, the cells were washed with ice-cold phosphate-buffered saline (PBS), resuspended in 100 μL PBS containing 5 μL of PE-labeled anti-CD11b (clone MEM-174, EXBIO Praha, Vestec, Czech Republic) or 5 μL of APC-labeled anti-CD14 (clone TÜK4), and incubated at room temperature in a light-protected area for 30 min. Then, the cells were washed with ice-cold PBS again and resuspended in 300 μL of PBS. All specimens were obtained by flow cytometry (FACSCalibur) and analyzed using FlowJo software (Tree Star). For morphology analysis, cells (1 × 10^5^) were adhered to microscopic slides using cytospin (Serocito, model 2400; FANEM, Guarulhos, Brazil), followed by Rosenfeld staining. The morphologies of the nucleus and cytoplasm of the treated cells were observed under a Leica DM2500 optical microscope, and images were acquired using LAS software version 4.6 (Leica, Bensheim, Germany).

### Statistical analysis

Statistical analysis was performed using GraphPad Instat 8 (GraphPad Software, Inc.). ANOVA and Bonferroni post-test and Student t test were used for comparisons, as appropriate. Correlograms were constructed using Morpheus. All *p* values were two-sided with a significance level of 5%.

## Results

### THZ-P1-2 reduces cell viability and induces DNA damage in leukemia cells

We investigated the role of PIP4Ks in the malignant phenotype of AML and ALL by using the candidate drug THZ-P1-2. Using a panel of 21 commonly used human myeloid and lymphoid leukemic cell lines, THZ-P1-2 caused a dose-dependent decrease in viability in these leukemia cells, with IC_50_ ranging from 5.6 to >100 µM for myeloid cells and 2.3 to 13.4 µM for lymphoid cells (Figure 1A). The reduction of cell viability induced by THZ-P1-2 was time-dependent in leukemic cells (Figure 1B). To get a better understanding of the cellular mechanisms causing the decrease in cell viability, we evaluated the DNA content of leukemia cells treated with vehicle or THZ-P1-2.

**Figure 1.**
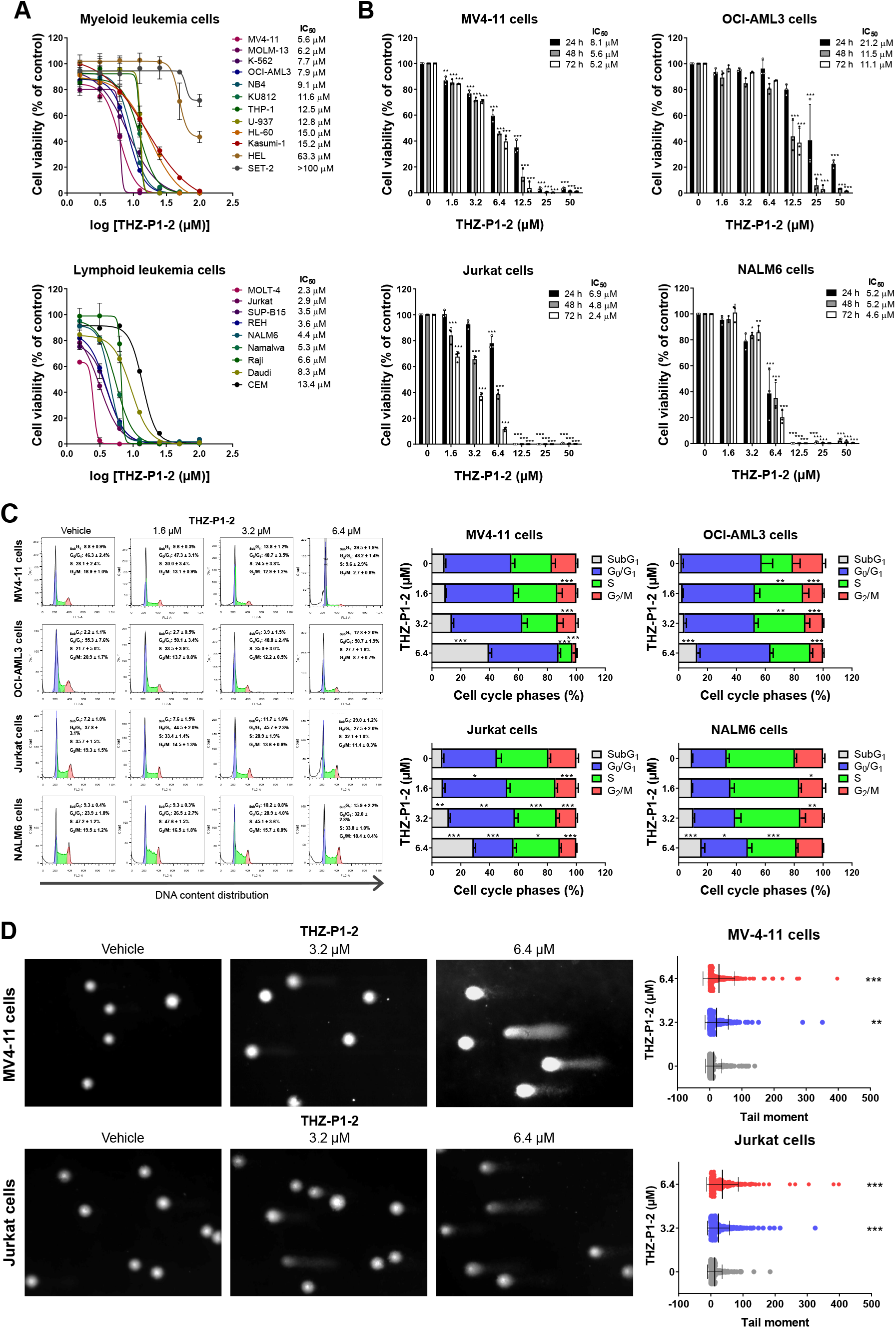
Pharmacological PIP4K2s inhibition reduces cell viability and induces DNA damage in leukemia cells. **(A)** Dose-response cytotoxicity was analyzed using a methylthiazoletetrazolium (MTT) assay in a panel of myeloid and lymphoid leukemic cell lines treated with vehicle or increasing concentrations of THZ-P1-2 (1.6, 3.2, 6.4, 12.5, 25, 50, and 100 μM) for 72 h. Values are expressed as the percentage of viable cells for each condition relative to vehicle-treated cells. The IC_50_ values and leukemia cell lines used are described in the Figure. **(B)** Dose- and time-response cytotoxicity was evaluated by methylthiazoletetrazolium (MTT) in MV4-11, OCI-AML3, Jurkat, and NALM6 cells treated with vehicle or with increasing concentrations of THZ-P1-2 (1.6, 3.2, 6.4, 12.5, 25, and 50 μM) for 24, 48, and 72 h. Bar graphs represent values expressed as a percentage for each condition relative to vehicle-treated controls. Results are presented as the mean ± SD of at least three independent experiments. The *p* values and cell lines are indicated in the graphs; * *p* < 0.05; ** *p* < 0.01; *** *p* < 0.001, ANOVA and Bonferroni post-test. **(C)** Phases of the cell cycle were determined by analyzing the DNA content by staining with propidium iodide and acquiring the data by flow cytometry following exposure of MV4-11, OCI-AML3, Jurkat, and NALM6 cells to the vehicle or THZ-P1-2 (1.6, 3.2, and 6.4 μM) for 24 h. A representative histogram for each condition is illustrated. The mean ± SD of at least three independent experiments is represented in the bar graph. The *p* values and cell lines are indicated in the graphs; * *p* < 0.05; ** *p* < 0.01; *** *p* < 0.001, ANOVA and Bonferroni post-test. **(D)** DNA damage was evaluated by the single-cell comet assay in MV4-11 and Jurkat treated with vehicle or THZ-P1-2 (3.2 or 6.4 µM) for 48 h. Scatter plots represent the tail moment values (head/tail DNA ratio) obtained using the LUCIA Comet Assay™ software. ** *p* < 0.01; *** *p* < 0.001, ANOVA and Bonferroni post-test.

Our results indicated cytostatic effects associated with a reduction of cells in S and G2/M phases when THZ-P1-2 was administrated at low concentrations (1.6 or 3.2 µM). Conversely, when THZ-P1-2 was administrated at high concentrations (6.4 µM), we observed cytotoxic effects linked with an increased accumulation of cells in the _sub_G1 (all *p* < 0.05, Figure 1C). The presence of DNA damage after exposure to THZ-P1-2 in leukemia cells was confirmed by the comet assay (all *p* < 0.05, Figure 1D).

### THZ-P1-2 triggers apoptosis and disrupts mitochondrial homeostasis and autophagic flux

Next, we evaluated the apoptotic effects of THZ-P1-2 in leukemic cells. An increase in apoptosis was detected in all cellular models upon the treatment with THZ-P1-2, whereby the highest apoptotic rate was observed in MV4-11 and Jurkat cells. (Figure 2A). In line with our apoptosis data, we also observed a significant loss of mitochondrial membrane potential (MMP) indicating mitochondrial damage (Figure 2B). However, the loss of MMP did not directly correlate with the frequency of apoptotic cells, especially in OCI-AML3 and NALM6 cells suggesting the activation of resistance/survival pathways.

**Figure 2.**
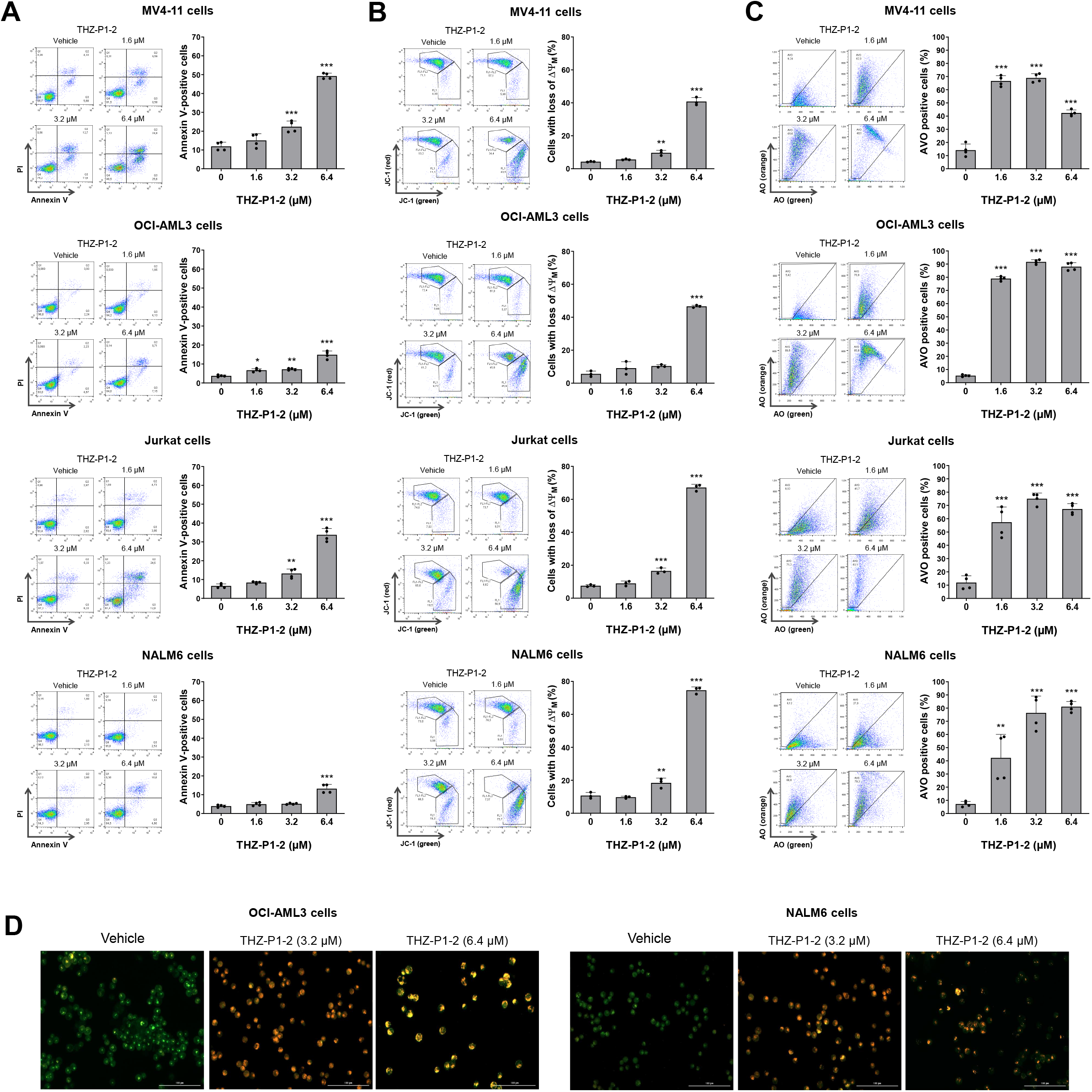
THZ-P1-2 induces apoptosis and dysfunction in mitochondria and autophagic flux. **(A)** Apoptosis was detected by flow cytometry in MV4-11, OCI-AML3, Jurkat, and NALM6 cells treated with vehicle or with increasing concentrations of THZ-P1-2 (1.6, 3.2, and 6.4 μM) for 24 h using an APC-annexin V/PI staining method. Representative dot plots are shown for each condition. The upper and lower right quadrants (Q2 plus Q3) cumulatively contain the apoptotic cell population (annexin V^+^ cells). Bar graphs represent the mean ± SD of at least three independent experiments. The *p* values and cell lines are indicated in the graphs; * *p* < 0.05, ** *p* < 0.01, *** *p* < 0.0001; ANOVA and Bonferroni post-test. **(B)** Mitochondrial membrane potential (ΔΨ_M_) analysis was evaluated using the JC-1 staining method and flow cytometry. Leukemic cells were treated with vehicle or THZ-P1-2 (1.6, 3.2, and 6.4 μM) for 24h. Representative dot plots are shown for each condition; the gate FL-2 contains cells with intact mitochondria and the gate FL-2/FL-1 contains cells with damaged mitochondria. Bar graphs represent the mean ± SD of at least three independent experiments and the p values are indicated; ** *p* < 0.01, *** *p* < 0.0001; ANOVA and Bonferroni post-test. **(C)** The evaluation of acidic vesicular organelles (AVOs) was investigated through acridine orange (AO) labeling and flow cytometry in AML and ALL cell lines treated with vehicle or THZ-P1-2 (1.6, 3.2, and 6.4 μM) for 24 h. Bar graphs represent the mean ± SD of at least three independent experiments and the p values are indicated; * *p* < 0.05, ** *p* < 0.01, *** *p* < 0.001; ANOVA and Bonferroni Post-test. **(D)** Alternatively, the presence of AVOs was confirmed by immunofluorescence on OCI-AML3 and NALM6 cell lines treated with vehicle or THZ-P1-2 (3.2 and 6.4 μM) for 72 h on a Lionheart FX automated microscope at magnification, ×400. The overlapping GFP and RFP channels are shown.

Previous studies, which used a murine genetic model deficient for *Pip4k2a*^-/-^ and *Pip4k2b*^-/-^ have shown that PIP4Ks are essential for the formation of autophagosomes [16]. Considering that autophagy has been linked to leukemogenesis as well as to the recycling of damaged mitochondria [31], we evaluated the formation of acidic vesicular organelles (AVO) as a mechanism of resistance. Notably, AVO levels were substantially elevated upon THZ-P1-2 exposure, particularly at concentrations that induced lower levels of apoptosis (Figure 2C-D), suggesting an inverse correlation between apoptosis and autophagy. At the molecular level, THZ-P1-2 induced the expression of apoptosis markers (PARP1 cleavage) and DNA damage (γH2AX) at levels compatible with those observed in previous cellular assays. Furthermore, LC3BII and SQSTM1/p62 levels were increased, which indicates a failure to complete the autophagic flow (Figure 3A).

**Figure 3.**
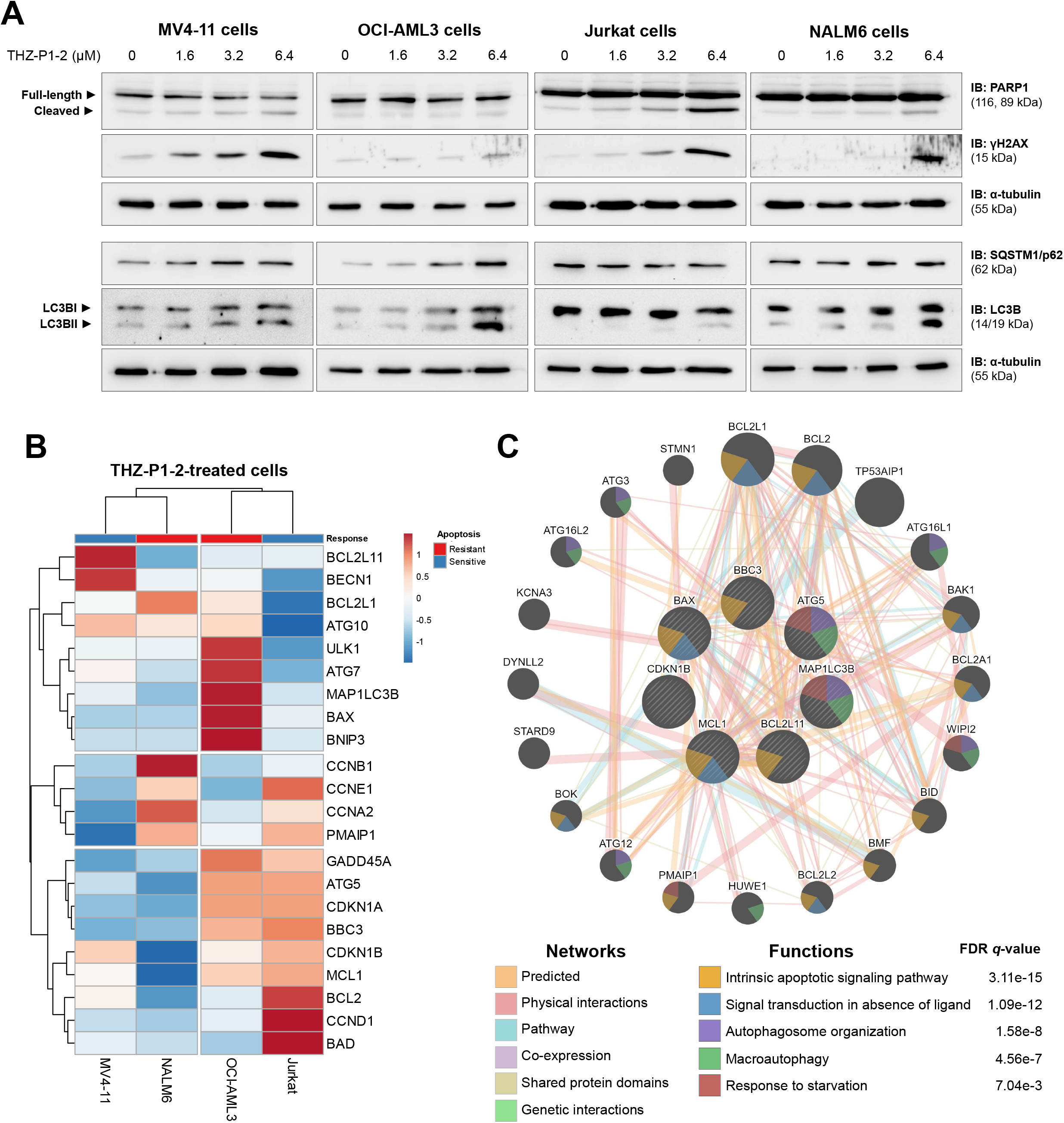
THZ-P1-2 induces markers of apoptosis, DNA damage, and blockage of autophagic flow. **(A)** Western blot analysis for PARP1, γH2AX, p62/SQSTM1, and LC3B in total extracts from MV4-11, OCI-AML3, Jurkat, and NALM6 cells treated with vehicle or with increasing doses of THZ-P1-2 (1.6, 3.2, and 6.4 µM) for 24 h. Membranes were re-incubated with α-tubulin antibody and developed with the SuperSignal™ West Dura Extended Duration Substrate system and GBox. **(B)** Heatmap depicting the gene expression profile of leukemic cell lines treated with vehicle or THZ-P1-2 (6.4 μM) for 24 h. The blue color in the heat map indicates decreased mRNA levels while red indicates induced mRNA levels, which were normalized by vehicle-treated cells (n = 4). **(C)** Network for THZ-P1-2 modulated genes constructed using the GeneMANIA database (https://genemania.org/). A total of seven genes (*BBC3, ATG5, MAP1LC3B, MCL1, CDKN1B, BAX*, and *BCL211*) were significantly modulated in all cell lines tested and are illustrated as crosshatched circles; the interacting genes included by modeling the software are indicated by circles without crosshatched. The main biological interactions and associated functions are described in the literature.

Exploratory analysis of genes associated with cell cycle progression, apoptosis, and autophagy, revealed that a total of seven genes were modulated in all leukemic models analyzed. These genes were significantly associated with the intrinsic apoptotic signaling pathway, signal transduction in absence of ligand, autophagosome organization, macroautophagy, and response to starvation (all FDR *q* < 0.05), corroborating the previous cellular and molecular findings (Figure 3B-C and Supplementary Table 5).

### THZ-P1-2 negatively impacts cellular metabolism in leukemia cells

Since the response to THZ-P1-2 was associated with mitochondrial damage and response to starvation in leukemic cells, we studied the effect of THZ-P1-2 on cellular metabolism. Our results confirmed that THZ-P1-2 exposure reduces the mitochondrial membrane potential of leukemia cells (Figure 4A). In addition, exposure to THZ-P1-2 reduced basal and maximal cellular respiration capacity (Figure 4B-D) as well as glycolytic flux (Figure 4E-F), indicating that the drug negatively impacts the metabolic state of leukemic cells.

**Figure 4.**
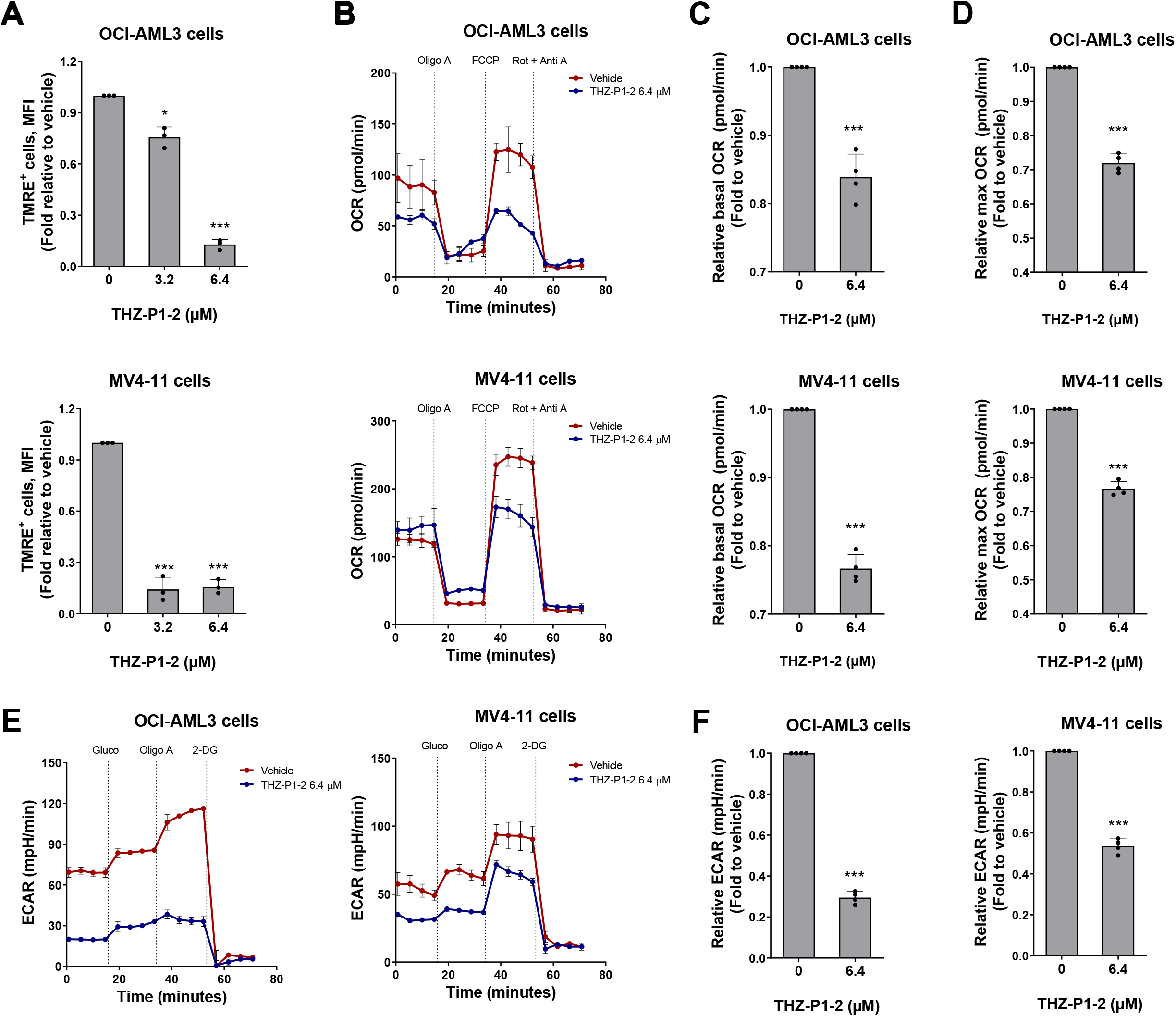
THZ-P1-2 reduces cellular metabolism in leukemic cells. **(A)** Mitochondrial membrane potential analysis was evaluated using the TMRE staining method and flow cytometry. OCI-AML3 and MV4-11 cells were treated with vehicle or THZ-P1-2 (3.2, and 6.4 μM) for 24 h. Values are expressed as the fold-change of vehicle-treated cells. Bar graphs represent the mean ± SD of at least three independent experiments and the p values are indicated; * *p* < 0.05, *** *p* < 0.0001; ANOVA and Bonferroni post-test. **(B)** Oxygen consumption rate (OCR) was determined in vehicle- or THZ-P1-2-treated (6.4 μM) OCI-AML3 and MV4-11 cells using a high-resolution respirometry. A representative line graph containing OCR upon sequential addition of oligomycin (Oligo A), carbonyl cyanide-p-trifluoromethoxyphenylhydrazone (FCCP), and rotenone plus antimycin (Rot + Anti A) are illustrated; OCR was measured over time. Values are expressed as the fold-change of vehicle-treated cells. Bar graphs represent the mean ± SD of OCR at baseline **(C)** and maximum state (upon FCCP) **(D)** of at least three independent experiments. The p values and cell lines are indicated in the graphs; *** *p* < 0.0001; Student t test. **(E)** Extracellular acidification rate (ECAR) was evaluated in vehicle- or THZ-P1-2-treated (6.4 μM) OCI-AML3 and MV4-11 cells by Seahorse XF96 analyzer. A representative line graph containing ECAR upon sequential addition of glucose (Gluco), oligomycin (Oligo A), and 2-deoxy-D-glucose (2-DG) is illustrated; ECAR was measured over time. **(F)** Values are expressed as the fold-change of vehicle-treated cells. Bar graphs represent the mean ± SD of at least three independent experiments. The p values and cell lines are indicated in the graphs; *** *p* < 0.0001; Student t test.

### THZ-P1-2 selectively reduces cell viability of primary leukemic blasts

To validate the antineoplastic potential of THZ-P1-2 in pre-clinical models of acute leukemias, primary cells from a heterogeneous set of patients diagnosed with *de novo* AML and ALL were submitted to *ex vivo* drug screenings. Similar to what was observed in the cell line model, THZ-P1-2 reduced the viability of primary leukemic cells in a dose-dependent manner with IC_50_ ranging from 6.4 to >100 µM for AML samples and 3.1 to 31.2 µM for ALL samples (Figure 5A). Furthermore, using a co-culture model system, similar results were observed in an independent and functionally well-characterized AML cohort (Figure 5B). Surprisingly, the response to THZ-P1-2 was correlated with a more metabolically active phenotype (oxygen consumption and extracellular acidification) of the leukemic blasts (Figure 5C). Among the genetic markers related to AML, the presence of mutations in *FLT3, NPM1, RUNX1, IDH1*/2, or *DNMT3A* were not associated with drug response (Figure 5D). In the context of healthy hematopoiesis, THZ-P1-2 only slightly reduced the cell proliferation of CB CD34^+^in long-term culture, while no difference in colony formation capacity was observed (Figure 5E-F).

**Figure 5.**
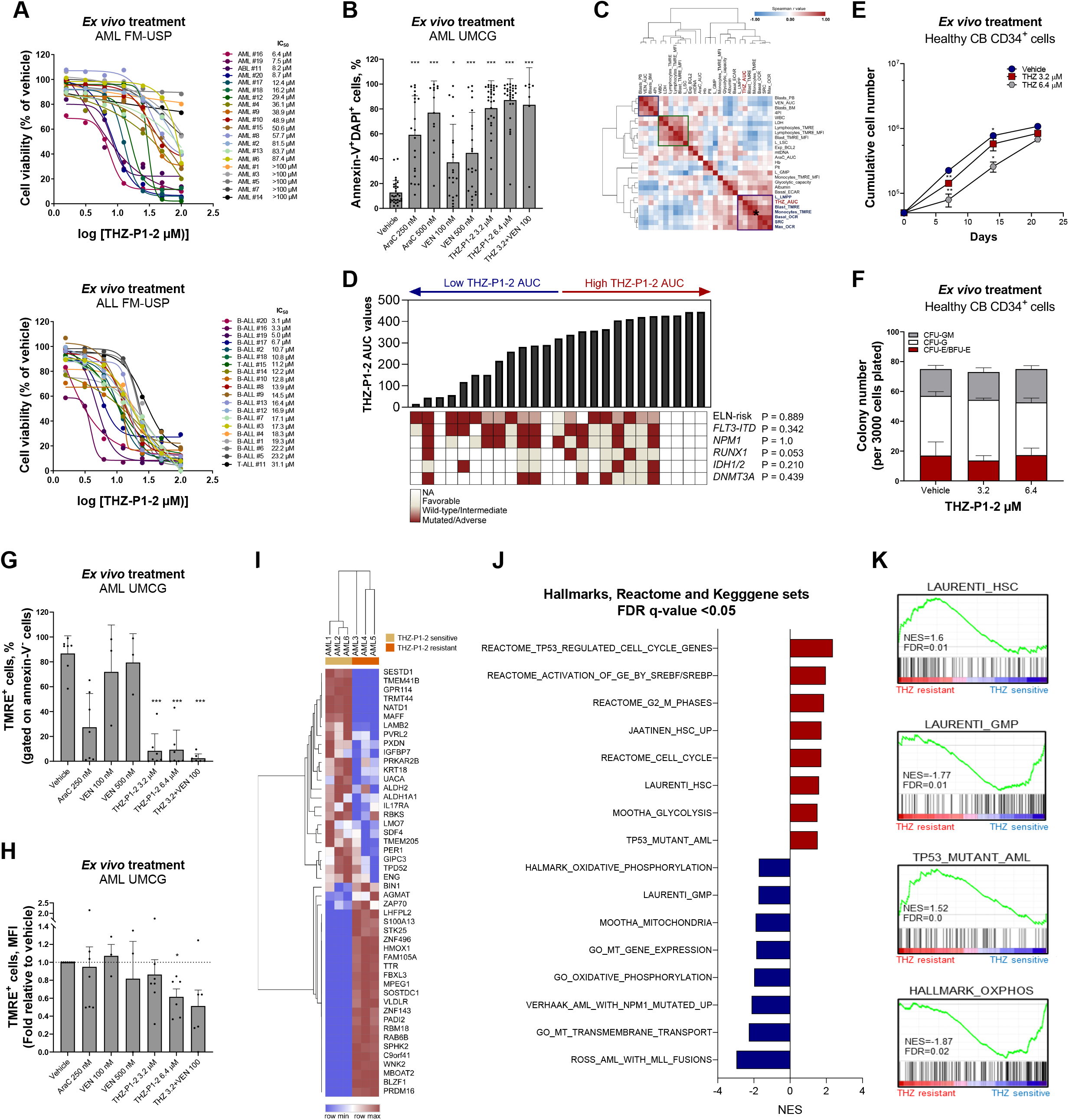
Pharmacological PIP4K2s inhibition selectively reduces cell viability of primary leukemic blasts. **(A)** Dose-response cytotoxicity was analyzed using a methylthiazoletetrazolium (MTT) assay in samples from acute myeloid leukemia (AML) or T- and B-acute lymphoblastic leukemia (T- or B-ALL) patients treated with vehicle or increasing concentrations of THZ-P1-2 (1.6, 3.2, 6.4, 12.5, 25, 50, and 100 μM) for 72 h. Values are expressed as the percentage of viable cells for each condition relative to vehicle-treated cells. The IC_50_ values for each patient sample are described. **(B)** Apoptosis was detected by flow cytometry in gated human Sca1^-^CD45^+^CD34^+^ (or CD117^+^ cells for *NPM1*-mutant AMLs) of acute myeloid leukemia (AML) samples in a co-culture system using a FITC-annexin V/DAPI staining method. Cells were treated with vehicle, cytarabine (AraC, 250 and 500 nM), venetoclax (VEN, 100 and 500 nM), and/or THZ-P1-2 (3.2 or 6.4 μM) for 72 h. Bar graphs represent the mean ± SD of all the independent patients screened, each point represents a patient. The *p* values are indicated in the graphs; * *p* < 0.05, ** *p* < 0.01, *** *p* < 0.0001; ANOVA and Bonferroni post-test. **(C)** Correlation analysis of THZ-P1-2 response with clinical, laboratory, molecular, and metabolic characteristics of the samples. Note that THZ-P1-2 responsiveness (area under curve, AUC) was clusterized with markers of mitochondrial metabolic markers (Levels of MMP in the blasts – Blast_TMRE; Basal OCR and Max OCR). Data were processed using the Morpheus platform (https://software.broadinstitute.org/morpheus/) **(D)** Association of THZ-P1-2 sensitivity (area under the curve, AUC), mutations in *FLT3, NPM1*, and *RUNX1*, and European LeukemiaNet (ELN) risk stratification. **(E)** Evaluation of long-term proliferation in neonatal cord blood (CB) CD34^+^ and primary AML mononuclear cells using a co-culture system. Data are expressed as mean ± SD of at least three independent experiments. The time points and *p* values are indicated in the graphs; * *p* < 0.05, ** *p* < 0.01, *** *p* < 0.0001; ANOVA and Bonferroni post-test. **(F)** Neonatal cord blood (CB) CD34^+^ cells were plated in cytokine-supplemented methylcellulose in the presence of vehicle or THZ-P1-2 (3.2 or 6.4 μM). Colonies were counted after 8-14 days of culture and are represented as the percent of vehicle-treated controls. Bars indicate the mean ± SD of at least three assays. **(G-H)** Mitochondrial membrane potential was detected by flow cytometry in gated human CD45^+^CD34^+^ (or CD117^+^ for *NPM1*-mutant AMLs)/annexin V^-^ from samples from acute myeloid leukemia (AML) in a co-culture system using the TMRE staining method. Cells were treated with vehicle, cytarabine (AraC, 250 nM), venetoclax (VEN, 100 or 500 nM), and/or THZ-P1-2 (3.2 [alone or in combination with VEN 100 nM] or 6.4 μM) for 72 h. Bar graphs represent the mean ± SD of at least three independent experiments, each point represents a patient. The *p* values are indicated in the graphs; * *p* < 0.05, ** *p* < 0.01, *** *p* < 0.0001; ANOVA and Bonferroni post-test. **(I)** Differently expressed proteins obtained from THZ-P1-2 (THZ)-sensitive (n = 3) and THZ-resistant (n = 3) AML patients were included in the heatmap (all false discovery rate (FDR) q-values (FDR q) < 0.25). **(J)** The bar graph represents the normalized enrichment scores (NES) for Hallmark, Reactome, and Kegg gene sets with FDR q < 0.05. **(K)** GSEA plots for enriched molecular signatures in THZ-P1-2 (THZ) resistant *vs*. sensitive AML patient’s proteome are also shown. NES and FDR q are indicated.

Mitochondrial dysfunction caused by THZ-P1-2 was also validated in primary leukemia cells (Figure 5G-H). Proteome analysis was performed on samples from patients considered sensitive and resistant to THZ-P1-2, which presented distinct protein signatures (Figure 5I). A gene enrichment analysis identified multiple processes and signaling pathways associated with mitochondrial metabolism, cell cycle stage, leukemic cell-of-origin, and TP53 pathway (all FDR q-value < 0.05, Figure 5J-K). Taken together, these results suggest that AML patients sensitive to THZ-P1-2 differ in their metabolic and proliferative state compared to resistant AMLs.

## Discussion

Here, we report the antileukemic effects of a novel PIP4K2 inhibitor, THZ-P1-2. In a targeted knockdown screen for phosphoinositide modulator-related genes, the *PIP4K2A* gene was identified as essential for the survival, proliferation, and clonogenicity of leukemia-initiating cells in humans and murine models [21]. High transcript levels of *PIP4K2A* and *PIP4K2C* were predictive of poor prognosis in AML patients [22,23]. Furthermore, SNPs in the regulatory region of the *PIP4K2A* gene were associated with increased mRNA levels and susceptibility to ALL development [19,20]. These studies revealed a potential role for this family of enzymes in the malignant phenotype of human leukemias that can serve as targets for treatment. Indeed, initial observations indicate that THZ-P1-2 reduces cell viability in six human leukemia cell lines, which was confirmed by us using a larger heterogeneous panel of leukemia cell lines and primary cells from two independent acute leukemia cohorts, but the mechanisms involved remain poorly explored until now [26].

In the present study, THZ-P1-2 reduced cell viability by disrupting mitochondrial homeostasis, cellular metabolism, and autophagy. The role of PIP4K2s in the regulation of energy metabolism and autophagy has been previously reported, in particular, the product of these enzymes, PI-4,5-P_2_, has been described as necessary for the fusion of lysosomes with autophagosomes and to complete the autophagic flux [16,18,26]. Considering our data, THZ-P1-2 was able to accurately phenocopy many of the cellular and molecular findings observed in PIP4K2s genetic inhibition models, including accumulation of SQSTM1/p62 and LC3BII [16]. There is also evidence that PIP4K2A suppresses lysosomal synthesis or enhances lysosomal turnover [16], which could explain the increase in AVOs upon treatment with THZ-P1-2 observed herein. In brief, our data indicate that THZ-P1-2 may exert a unique pharmacological activity, acting as an inducer of the initial stages of autophagy through mitochondrial dysfunction and as an inhibitor of the final stages of autophagy. Thus, THZ-P1-2 may leave leukemia cells in a state of “autophagic catastrophe”, which reduces cell viability, even in the absence of classical cell death pathways (*i*.*e*. apoptosis). Impairment of autophagy also promotes the accumulation of mitochondrial reactive oxygen species and DNA damage [31], which was confirmed by our data.

The reduction in cellular metabolism observed upon THZ-P1-2 exposure was timely in the context of acute leukemias. Corroborating our findings, it has been demonstrated that loss of the PIP4K2s leads to alterations in mitochondrial structural and functional integrity, as observed in double PIP4K2A/B knockout mice or inhibition by shRNA in solid tumor cell lines [25]. Recent data have indicated that leukemia cells may rely more on mitochondrial metabolism, being more responsive to venetoclax [8,32]. Since THZ-P1-2 led to mitochondrial damage and reduced cellular metabolism, we investigated the effects of the combination of both drugs on OCI-AML3 cells, a leukemia cell line highly resistant to venetoclax. The combined treatment showed synergistic effects, reducing the IC_50_ of venetoclax and strongly increasing the induction of apoptosis (Supplementary Figure 1). Clinical trials with venetoclax-based regimens have shown encouraging results in hematological malignancies (*i*.*e*. AML) and the search for drug resistance markers as well as mechanisms to overcome them has been a very active field of research [33,34]. Our preliminary findings suggest THZ-P1-2 as a potential candidate in this context but require further investigation. Furthermore, OXPHOS-inhibition has been associated with disruption of stemness properties in acute leukemia [7,35,36], which prompted us to evaluate the effects of THZ-P1-2 on cell differentiation of myeloid models. Drug treatment up-regulated the expression of differentiation markers and induced morphological changes compatible with a more differentiated phenotype (Supplementary Figure 2). Our results are in line with a recent study suggesting PIP4K2s be involved in the regulation of organelle crosstalk as well as the preservation of metabolic homeostasis [18]. The minimal effects observed in healthy CB CD34^+^ cells suggest that THZ-P1-2 may have a favorable therapeutic window. Indeed, PIP4K2A depletion attenuated the growth of primary leukemia blasts but did not significantly impact the clonogenic or multilineage differentiation potential of healthy hematopoietic stem cells [21].

Our proteome analysis revealed that THZ-P1-2 resistant AML CD34^+^ (or CD117^+^ for *NPM1*-mut AMLs) cells display a molecular signature similar to L-HSC and TP53 mutant AML, with a more glycolytic-driven metabolism, whereas THZ-P1-2 sensitive AML samples display enrichment for cell maturation markers and mitochondrial metabolism, which corroborates the findings of our functional studies. The role of TP53 in the context of PIP4K2s has been recognized in several studies. In *TP53*-mutated cancer cells, the inhibition of PIP4K2A/B reduced cell survival. Similar results were observed in TP53^−/−^ PIP4K2A^−/−^ PIP4K2B^+/−^ mice, which presented a lower tumor burden and increased tumor-free survival[24]. Lundquist *et al*. [16] showed that *TP53* silencing enhances the role of PIP4K2A/B inhibition in murine cell autophagy, reinforcing TP53 as an important player in the response to PIP4Ks inhibition.

In conclusion, the pharmacological PIP4Ks inhibitor, THZ-P1-2, disrupts mitochondrial metabolism and autophagic flux, two cellular processes that are being proposed as promising targets to improve therapeutic responses in acute leukemia. Subsequently, our data shed light on a new class of drugs to increase the range of treatment options for leukemia.

## Author contributions

Conceptualization: K.L., E.M.R, J.A.M-N; investigation: K.L., D.A.P.M., L.B.L.M., I.W., J.J.S., E.G., E.M.R., J.A.M-N.; technical assistance and discussion: J.L.C-S., G.S.L., R.C.C., A.M.L., W.F.S.J., A.P.A.L.L., E.D.R.P.V., J.R.H., G.H.; resources: G.H.; data curation: K.L., D.A.P.M., J.J.S., E.M.R, J.A.M-N; writing—original draft: K.L., D.A.P.M., J.J.S., E.M.R, J.A.M-N; writing—review and editing: L.B.L.M., J.L.C-S., G.S.L., I.W., R.C.C., A.M.L., W.F.S.J., A.P.A.L.L., E.D.R.P.V., E.G., J.R.H., G.H.; funding acquisition: J.J.S. E.M.R, and J.A.M-N.; overall supervision: E.M.R. and J.J.S

## Acknowledgements

K.L. received a fellowship from FAPESP (Grant #2020/12842-0). I.W. received a fellowship from FAPESP (Grant #2015/09228-0). D.A.P-M. received a fellowship from FAPESP (Grant #2017/23117-1). I.W and D.A.P-M were sponsored by the Abel Tasman Talent Program (ATTP) of the Graduate School of Medical Sciences of the University of Groningen/University Medical Center Groningen (UG/UMCG), The Netherlands. This study was supported by grant #2019/23864-7 and #2021/11606-3 from the São Paulo Research Foundation (FAPESP). This study was financed in part by the Coordenação de Aperfeiçoamento de Pessoal de Nível Superior - Brasil (CAPES) - Finance Code 001. The authors thank Prof. Carlos Frederico Martins Menck for providing assistance with the comet assay and critical review of the manuscript.

## Data availability

All data are also available from the corresponding author on request. LFQ proteome data are deposited under PRIDE PXD030463. The remaining data are available in the Article and Supplementary Information. Source data are provided in this paper.

## Figure legends

**Supplementary Figure 1. THZ-P1-2 potentates venetoclax-induced apoptosis in OCI-AML3 cells. (A)** Dose-response cytotoxicity for combined treatment was analyzed by methylthiazoletetrazolium (MTT) assay for OCI-AML3 cells treated with graded concentrations of venetoclax and THZ-P1-2 alone or in combination with each other for 48 h, as indicated. Values are expressed as the percentage of viable cells for each condition relative to vehicle-treated cells. Results are shown as the mean of at least three independent experiments. Note that the inhibitory concentration of 50% (IC_50_) for venetoclax was reduced in combination with THZ-P1-2 in OCI-AML3 cells. **(B)** Apoptosis was detected by flow cytometry in OCI-AML3 cells treated with venetoclax and/or THZ-P1-2 for 48 hours using an APC-annexin V/PI staining method. Representative dot plots are shown for each condition; the upper and lower right quadrants (Q2 plus Q3) cumulatively contain the apoptotic population (annexin V^+^ cells). Bar graphs represent the mean ± SD of at least three independent experiments. The p values and cell lines are indicated in the graphs; \**p* < 0.05 for venetoclax- and/or THZ-P1-2-treated cells vs. vehicle-treated cells, ^#^*p* < 0.05 for venetoclax- or THZ-P1-2-treated cells *versus* combination treatment at the corresponding doses; ANOVA and Bonferroni post-test.

**Supplementary Figure 2. THZ-P1-2 induces cell differentiation in acute myeloid leukemia cellular models**. OCI-AML3, THP-1, and NB4 cells were exposed to vehicle or THZ-P1-2 (1.6, 3.2, and 6.4 μM) for 72 h. The histogram represents the mean fluorescence intensity (M.F.I.) for PE-CD11b **(A)** and APC-CD14 **(B)** expression. Bar graphs represent the mean ± SD of at least three independent experiments. The *p* values are indicated; \**p* < 0.05, \*\**p* < 0.01, \*\*\**p* < 0.001; ANOVA and Bonferroni post-test. **(C)** Cytospin preparations were stained with Rosenfeld. The cell lines and THZ-P1-2 concentrations are indicated in the images. Scale bar = 50 μm.

