## Supplementary Material for "The PIP4K2 inhibitor THZ-P1-2 exhibits antileukemia activity by disruption of mitochondrial homeostasis and autophagy"

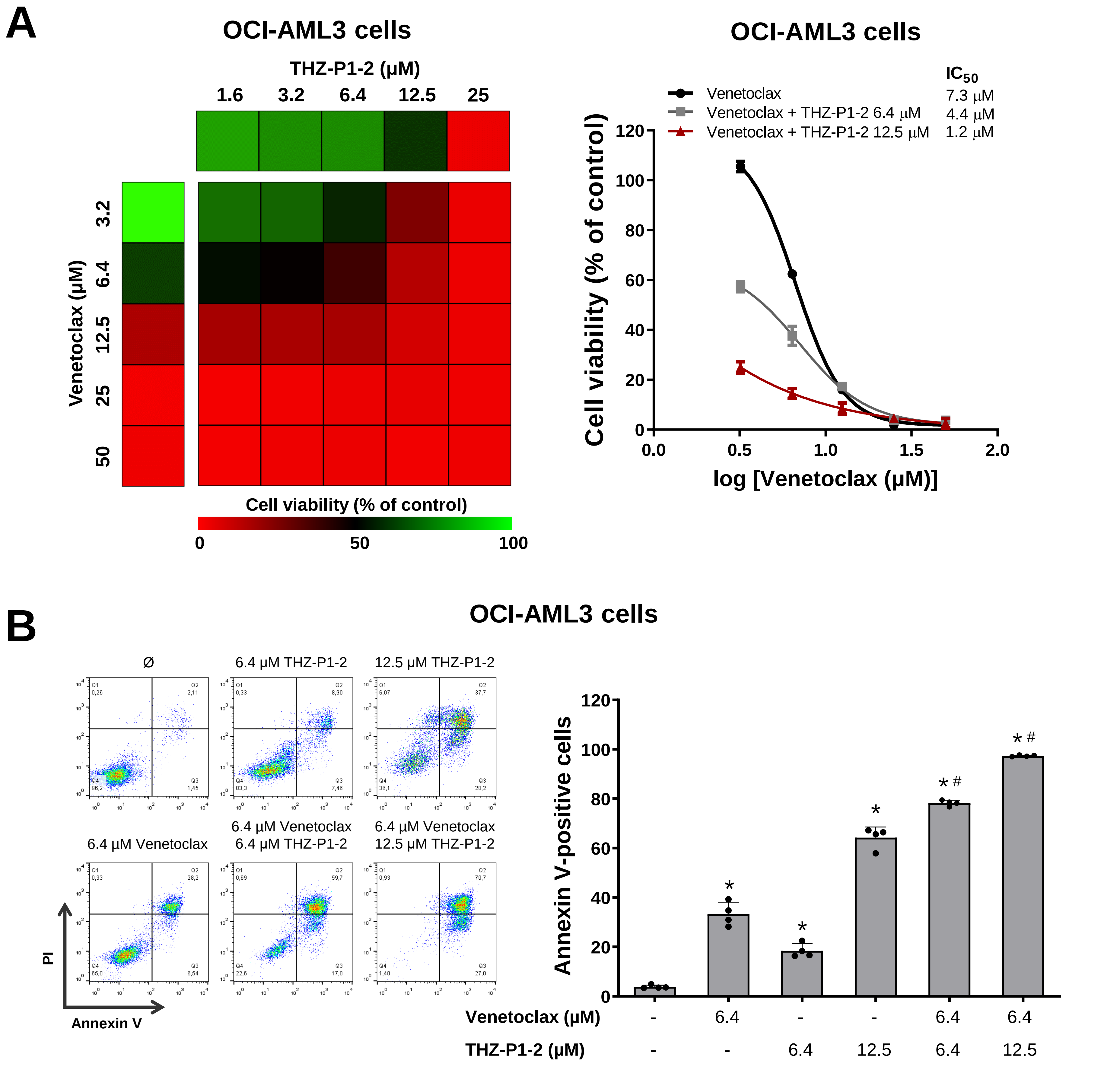
A
OCI-AML3 cells
THZ-P1-2 (μM)
1.6
3.2
6.4
12.5
25
Venetoclax (μM)
3.2
6.4
12.5
25
50
Cell viability (% of control)
0
50
100
OCI-AML3 cells
IC50
Venetoclax 7.3 μM
Venetoclax + THZ-P1-2 6.4 μM 4.4 μM
Venetoclax + THZ-P1-2 12.5 μM 1.2 μM
Cell viability (% of control)
log [Venetoclax (μM)]
B
OCI-AML3 cells
Ø
6.4 μM THZ-P1-2
12.5 μM THZ-P1-2
6.4 μM Venetoclax
6.4 μM Venetoclax 6.4 μM THZ-P1-2
6.4 μM Venetoclax 12.5 μM THZ-P1-2
PI
Annexin V
Annexin V-positive cells
Venetoclax (μM) - 6.4 - - 6.4 6.4
THZ-P1-2 (μM) - - 6.4 12.5 6.4 12.5

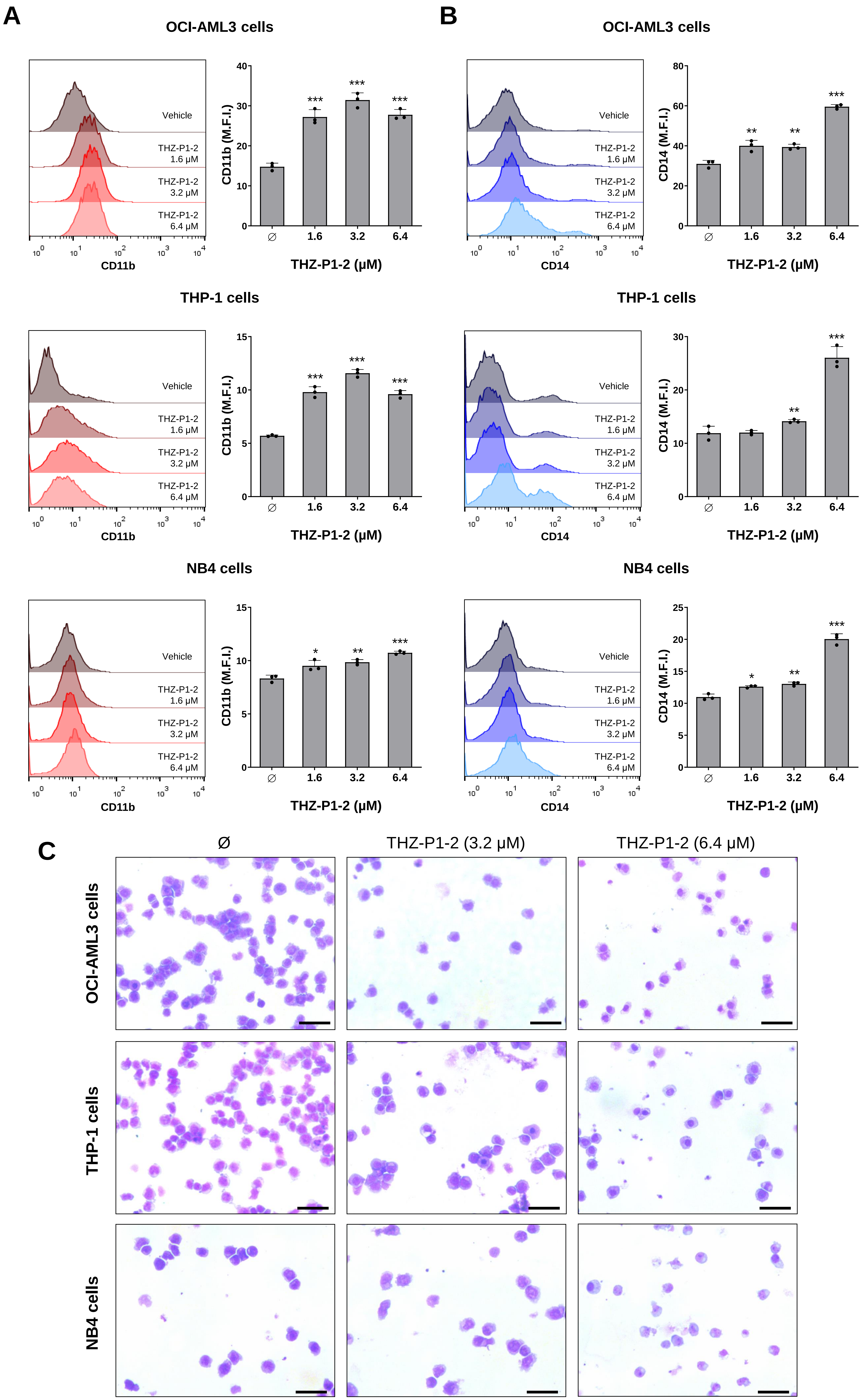

A
B
C
OCI-AML3 cells
THP-1 cells
NB4 cells
Vehicle
THZ-P1-2 1.6 μM
THZ-P1-2 3.2 μM
THZ-P1-2 6.4 μM
CD11b
CD14
CD11b (M.F.I.)
CD14 (M.F.I.)
THZ-P1-2 (μM)
Ø
1.6
3.2
6.4
Ø
THZ-P1-2 (3.2 μM)
THZ-P1-2 (6.4 μM)

| MTT code | THZ-P1-2 IC50 (uM) | Sex | Blasts (%) | Disease | Karyotype | t(15;17) PML::RARA | t(8;21) RUNX1::RUNX1T1 | inv(16) CBFB::YH11 | MLL::AF4 |
|---|---|---|---|---|---|---|---|---|---|
| AML #1 | >100 | Male | 61.6 | AML with recurrent genetic abnormalities | 46,Y,t(X;1)(q26;p36.1)[7]/46,idem,inv(16)(p13q22)[13] | NA | non-mutant | mutant | NA |
| AML #2 | 81.5 | Female | 82.4 | Acute Myeloid Leukemia Not Otherwise Specified | 47,XX,t(3;12)(q26;q14),del(5)(q13q31),-7,+8,+mar[11]/46,XX[4] | NA | NA | NA | NA |
| AML #3 | >100 | Female | 69.4 | AML with recurrent genetic abnormalities | 46,XX | NA | non-mutant | non-mutant | NA |
| AML #4 | 36.1 | Male | 35.6 | Acute myeloid leukaemia with myelodysplasia-related changes | 47,XY,?del(5)(q?13q?31),-18,+2mar | NA | non-mutant | non-mutant | NA |
| AML #5 | >100 | Female | 22.0 | Acute Myeloid Leukemia Not Otherwise Specified | 46,XX | non-mutant | non-mutant | non-mutant | NA |
| AML #6 | 87.4 | Female | 86.4 | AML with recurrent genetic abnormalities | 46,XX | NA | non-mutant | non-mutant | NA |
| AML #7 | >100 | Male | 27.6 | Acute Myeloid Leukemia Not Otherwise Specified | 47,XY,+11[2]/46,XY[18] | NA | non-mutant | non-mutant | NA |
| AML #8 | 57.7 | Female | 69.5 | Acute Myeloid Leukemia Not Otherwise Specified | NA | non-mutant | non-mutant | non-mutant | NA |
| AML #9 | 38.9 | Female | 86 | Acute Myeloid Leukemia Not Otherwise Specified | 46,XX | NA | non-mutant | non-mutant | NA |
| AML #10 | 48.9 | Male | 25 | Acute Myeloid Leukemia Not Otherwise Specified | 46,XY | NA | non-mutant | non-mutant | NA |
| AML #11 | 8.2 | Female | NA | Biphenotypic acute leukaemia | NA | NA | NA | NA | NA |
| AML #12 | 29.4 | Male | 53.6 | AML with recurrent genetic abnormalities | 45,X,-Y,t(8;21)(q22;q22)[12]/46,XY[8] | NA | mutant | non-mutant | NA |
| AML #13 | 83.7 | Male | 80 | Acute Myeloid Leukemia Not Otherwise Specified | 46,XY,-5,+8,+8,der(10)t(8;10)(q11.2;p15),-17[20] | NA | non-mutant | non-mutant | NA |
| AML #14 | >100 | Male | 95 | AML with recurrent genetic abnormalities | 46,XY | NA | NA | NA | NA |
| AML #15 | 50.6 | Female | 84 | Acute Myeloid Leukemia Not Otherwise Specified | 46,XX | non-mutant | non-mutant | non-mutant | NA |
| AML #16 | 6.4 | Female | 66.8 | Acute Myeloid Leukemia Not Otherwise Specified | NA | NA | non-mutant | NA | NA |
| AML #17 | 12.4 | Female | 86.4 | Acute Myeloid Leukemia Not Otherwise Specified | 46,XX | NA | non-mutant | non-mutant | NA |
| AML #18 | 16.2 | Male | 91.6 | NA | 46,XY | NA | non-mutant | non-mutant | NA |
| AML #19 | 7.5 | Female | 89.6 | AML with recurrent genetic abnormalities | 46,XX | non-mutant | non-mutant | non-mutant | NA |
| AML #20 | 8.7 | Female | 96 | AML with recurrent genetic abnormalities | NA | mutant | non-mutant | non-mutant | NA |

| MTT code | NPM1 | FLT3-ITD | FLT3-TKD | KIT | CEBPA | DNMT3A | BCR::ABL1 (P190) | BCR::ABL1 (P210) | TELL::AML1 | ASXL1 | IDH1 | IDH2 |
|---|---|---|---|---|---|---|---|---|---|---|---|---|
| AML #1 | non-mutant | non-mutant | mutant | non-mutant | non-mutant | NA | non-mutant | non-mutant | NA | NA | non-mutant | NA |
| AML #2 | non-mutant | non-mutant | non-mutant | non-mutant | non-mutant | NA | NA | NA | NA | NA | non-mutant | mutant |
| AML #3 | non-mutant | mutant | non-mutant | non-mutant | non-mutant | NA | non-mutant | non-mutant | NA | non-mutant | non-mutant | non-mutant |
| AML #4 | non-mutant | non-mutant | non-mutant | non-mutant | non-mutant | NA | non-mutant | non-mutant | non-mutant | non-mutant | non-mutant | non-mutant |
| AML #5 | non-mutant | non-mutant | non-mutant | non-mutant | non-mutant | NA | non-mutant | NA | NA | NA | NA | NA |
| AML #6 | mutant | non-mutant | non-mutant | non-mutant | mutant | mutant | non-mutant | non-mutant | NA | mutant | non-mutant | non-mutant |
| AML #7 | non-mutant | non-mutant | non-mutant | NA | non-mutant | NA | non-mutant | non-mutant | NA | NA | NA | NA |
| AML #8 | non-mutant | mutant | non-mutant | non-mutant | non-mutant | non-mutant | non-mutant | non-mutant | non-mutant | non-mutant | non-mutant | non-mutant |
| AML #9 | mutant | non-mutant | mutant | NA | non-mutant | NA | non-mutant | non-mutant | NA | non-mutant | non-mutant | non-mutant |
| AML #10 | non-mutant | non-mutant | non-mutant | non-mutant | non-mutant | NA | non-mutant | non-mutant | non-mutant | non-mutant | non-mutant | non-mutant |
| AML #11 | NA | NA | NA | NA | NA | NA | NA | NA | NA | NA | NA | NA |
| AML #12 | non-mutant | non-mutant | mutant | non-mutant | non-mutant | NA | NA | non-mutant | NA | non-mutant | non-mutant | non-mutant |
| AML #13 | non-mutant | non-mutant | non-mutant | NA | non-mutant | NA | NA | non-mutant | NA | non-mutant | non-mutant | mutant |
| AML #14 | mutant | mutant | non-mutant | NA | NA | NA | NA | non-mutant | NA | NA | NA | NA |
| AML #15 | non-mutant | non-mutant | non-mutant | NA | non-mutant | NA | NA | NA | NA | NA | NA | NA |
| AML #16 | mutant | mutant | non-mutant | NA | non-mutant | mutant | non-mutant | non-mutant | NA | NA | NA | NA |
| AML #17 | mutant | non-mutant | mutant | NA | mutant | mutant | non-mutant | non-mutant | NA | NA | NA | NA |
| AML #18 | non-mutant | non-mutant | non-mutant | non-mutant | non-mutant | mutant | non-mutant | non-mutant | NA | mutant | non-mutant | mutant |
| AML #19 | mutant | non-mutant | non-mutant | non-mutant | non-mutant | non-mutant | non-mutant | non-mutant | NA | non-mutant | mutant | non-mutant |
| AML #20 | non-mutant | mutant | non-mutant | NA | non-mutant | non-mutant | non-mutant | non-mutant | NA | NA | NA | NA |

| MTT code | THZ-P1-2 IC50 (uM) | Sex | Blasts (%) | Disease | CRLF2 rearrangement | TEL::AML1 | MLL::AF4 | TCF3::PBX1/(1;19) | MLPA-IKA (IKAROS) | Karyotype |
|---|---|---|---|---|---|---|---|---|---|---|
| ALL #1 | 19.3 | Male | 94.4 | B Precursor Cell Leukemia | mutant | NA | non-mutant | non-mutant | NA | 46,XY |
| ALL #2 | 10.7 | Male | 78,0 | B Precursor Cell Leukemia | non-mutant | NA | NA | NA | NA | 46,XY |
| ALL #3 | 17.3 | Male | 26,0 | B Precursor Cell Leukemia | NA | NA | NA | NA | NA | 46,XY |
| ALL #4 | 18.3 | Female | 90.4 | B Precursor Cell Leukemia | NA | non-mutant | non-mutant | non-mutant | Deleted | 46,XX |
| ALL #5 | 23.2 | Female | 76.4 | B Precursor Cell Leukemia | mutant | NA | non-mutant | non-mutant | Deleted | 46,XX,inv(9)(p13q34)[18]/46,XX[2] |
| ALL #6 | 22.2 | Male | 97.2 | B Precursor Cell Leukemia | mutant | non-mutant | non-mutant | non-mutant | Deleted | 46,XY |
| ALL #7 | 17.1 | Male | 77.4 | B Precursor Cell Leukemia | NA | non-mutant | non-mutant | non-mutant | NA | 46,XY,t(9;22)(q34;q11.2)[7]/46,XY[2] |
| ALL #8 | 13.9 | Female | 90.4 | B Precursor Cell Leukemia | mutant | NA | non-mutant | non-mutant | NA | 46,XX |
| ALL #9 | 14.5 | Male | 91.2 | B Precursor Cell Leukemia | mutant | non-mutant | non-mutant | non-mutant | Deleted | 46,XY |
| ALL #10 | 12.8 | Female | 90.4 | B Precursor Cell Leukemia | non-mutant | NA | non-mutant | non-mutant | NA | NA |
| ALL #11 | 31.2 | Male | 69.2 | Leukemia/Lymphoblastic T lymphoma | NA | NA | non-mutant | non-mutant | Deleted | 47,XY,add(1)(p36.1),add(2)(p23),del(5)(q31),del(6)(q21q25-27),+8,del(12)(p12),del(20)(q11.2)[4]/46,XY[16] |
| ALL #12 | 16.9 | Female | 92.8 | B Precursor Cell Leukemia | mutant | non-mutant | non-mutant | non-mutant | Deleted | 46,XX |
| ALL #13 | 16.4 | Male | 96.4 | B Precursor Cell Leukemia | non-mutant | non-mutant | non-mutant | non-mutant | NA | NA |
| ALL #14 | 12.2 | Male | 88,0 | B Precursor Cell Leukemia | NA | NA | NA | NA | Deleted | NA |
| ALL #15 | 11.2 | Male | 92,0 | Mature T cell neoplasm | NA | NA | NA | NA | NA | 44,XY,-6,-8,add(11)(q13), der(14)t(8;14)(q11.2;p11.2)x2,-21,+mar[19]/46,XY[1] |
| ALL #16 | 3.3 | Male | 42,0 | B Precursor Cell Leukemia | non-mutant | NA | non-mutant | non-mutant | Deleted | 50,XY,+add(1)(p22),-2,+add(3)(q12),+6,del(6)(q13q23)x2.+10,del(14)(q24),+mar[14]/46,XY[6] |
| ALL #17 | 6.7 | Female | 94,0 | B Precursor Cell Leukemia | NA | NA | NA | NA | NA | 46,XX |
| ALL #18 | 10.8 | Male | 87,0 | B Precursor Cell Leukemia | mutant | non-mutant | non-mutant | non-mutant | Deleted | 46,XY |
| ALL #19 | 5.0 | Male | 78.6 | B Precursor Cell Leukemia | NA | NA | NA | NA | NA | NA |
| ALL #20 | 3.1 | Female | 69.2 | B Precursor Cell Leukemia | NA | non-mutant | non-mutant | non-mutant | Deleted | NA |

| AML_samples | THZ-P1-2 AUC | ELN-risk stratification | Karyotype | Sex | Disease | RUNX1 | Spliceosome_genes | DNMT3A | RAS/PTPN11 | FLT3-ITD | NPM1 | CEBPA | IDH1_2 |
|---|---|---|---|---|---|---|---|---|---|---|---|---|---|
| #1 | 16 | Adverse | 45,XX, +3p, -7, -8p, 46,XX | Female | AML post MDS | NA | NA | NA | NA | non-mutant | NA | NA | NA |
| #2 | 405,4 | Adverse | 46,XY | Male | AML | mutant | non-mutant | mutant | non-mutant | mutant | mutant | non-mutant | non-mutant |
| #3 | 338,2 | NA | NA | NA | atypical CML | NA | NA | NA | NA | NA | NA | NA | NA |
| #4 | 290,6 | Adverse | 46,XY, t(7;11) | Male | AML | NA | NA | NA | NA | mutant | non-mutant | NA | NA |
| #5 | 259 | Adverse | 46,XX | Female | mixed phenotype Acute leukemia | NA | NA | NA | NA | mutant | NA | NA | mutant |
| #6 | 117,8 | Adverse | complex karyotype | Female | mixed phenotype Acute leukemia | NA | NA | NA | NA | NA | non-mutant | NA | NA |
| #7 | 56,16 | Intermediate | 46,XY | Male | AML | NA | NA | NA | NA | mutant | mutant | NA | NA |
| #8 | 443,8 | Intermediate | 46,XX | Female | AML | NA | NA | NA | NA | mutant | mutant | NA | NA |
| #9 | 354,9 | Adverse | 46,XX,7q- | Female | AML | NA | NA | NA | NA | non-mutant | non-mutant | NA | NA |
| #10 | 364,2 | Intermediate | 46,XX | Female | AML | non-mutant | non-mutant | mutant | non-mutant | mutant | mutant | non-mutant | non-mutant |
| #11 | 321,4 | Intermediate | 46,XX | Female | AML | non-mutant | non-mutant | mutant | non-mutant | mutant | mutant | non-mutant | non-mutant |
| #12 | 288 | NA | NA | NA | AML | NA | NA | NA | NA | NA | NA | NA | NA |
| #13 | 281,6 | Favorable | 46,XX | Female | AML | NA | NA | NA | NA | non-mutant | mutant | non-mutant | NA |
| #14 | 445,6 | Adverse | 46,XX | Female | AML | mutant | NA | NA | NA | mutant | non-mutant | non-mutant | NA |
| #15 | 427,2 | NA | NA | NA | AML | non-mutant | non-mutant | mutant | non-mutant | mutant | mutant | non-mutant | non-mutant |
| #16 | 410,9 | Adverse | 45,XX,add(3)(q21),der(5)?t(5;7)(q15;p21),-7,add(11)(q2?2),add(12)(p12),add(17)(q23) [13]/45,idem,?add(9)(q32)[4]/46,XX[3]. | Female | AML from MDS/CLL | NA | NA | NA | NA | non-mutant | non-mutant | non-mutant | NA |
| #17 | 151,4 | Adverse | 45,XY,-7,del(12)(p11p12) | Male | sAML | non-mutant | mutant | mutant | mutant | non-mutant | non-mutant | non-mutant | non-mutant |
| #18 | 43,9 | Intermediate | 46,XY | Male | AML | non-mutant | non-mutant | non-mutant | non-mutant | mutant | mutant | non-mutant | mutant |
| #19 | 425,3 | Adverse | 46,XY,?inv(2)(p?21q?21),del(5)(p13p15),-7[10] | Male | AML | mutant | non-mutant | non-mutant | non-mutant | non-mutant | non-mutant | non-mutant | non-mutant |
| #20 | 151,5 | Intermediate | 46,XY | Male | AML | non-mutant | mutant | non-mutant | non-mutant | non-mutant | non-mutant | non-mutant | non-mutant |
| #21 | 357,4 | Intermediate | 46,XY | Male | AML | non-mutant | non-mutant | mutant | non-mutant | mutant | mutant | non-mutant | mutant |
| #22 | 420,4 | NA | NA | Male | AML | NA | NA | NA | NA | NA | NA | NA | NA |
| #23 | 216 | NA | NA | Female | AML | NA | NA | NA | NA | NA | NA | NA | NA |
| #24 | 47,36 | NA | NA | NA | AML | NA | NA | NA | NA | NA | NA | NA | NA |
| #25 | 429 | NA | NA | NA | AML | NA | NA | NA | NA | NA | NA | NA | NA |

**Supplementary Table 4.** Primer sequences and concentrations.

| Gene | Sequence | Concentration |
|---|---|---|
| *ATG5* | FW: GGGCCATCAATCGGAAAC<br>RV: AGCCACAGGACGAAACAG | 300 nM |
| *ATG7* | FW: CGTTGCCCACAGCATCATCTTC<br>RV: TCCCATGCCTCCTTTCTGGTTC | 300 nM |
| *ATG10* | FW: TACGCAACAGGAACATCCA<br>RV: AACAACTGGCCCTACAATGC | 150 nM |
| *BAD* | FW: CACCAGCAGGAGCAGCCAAC<br>RV: CGACTCCGGATCTCCACAGC | 300 nM |
| *BAX* | FW: GAGCTGCAGAGGATGATTGC<br>RV: CAGCTGCCACTCGGAAAA | 300 nM |
| *BBC3* | FW: GACCTCAACGCACAGTACGAG<br>RV: AGGAGTCCCATGATGAGATTG | 300 nM |
| *BCL2* | FW: ATGTGTGTGGAGAGCGTCAA<br>RV: ACAGTTCCACAAAGGCATCC | 300 nM |
| *BCL2L11* | FW: ATGTCTGACTCTGACTCTCG<br>RV: CCTTGTGGCTCTGTCTGTAG | 300 nM |
| *BECN1* | FW: TCTGAAGAGGACCTGGACCCT<br>RV: GGCTCACGTCCATCTCGTC | 300 nM |
| *BNIP3* | FW: ATATGGGATTGGTCAAGTCGG<br>RV: CGCTCGTGTTCCTCATGCT | 300 nM |
| *CCNA2* | FW: GCCTTTCATTTAGCACTCTACA<br>RV: CAGGGTATATCCAGTCTTTCG | 300 nM |
| *CCNB1* | FW: GTCTCCATTATTGATCGGTTCATG<br>RV: CCAATTTCTGGAGGGTACATTTCT | 300 nM |
| *CCND1* | FW: CTCGGTGTCCTACTTCAAATG<br>RV: AGCGGTCCAGGTAGTTCAT | 300 nM |
| *CCNE1* | FW: TATATGGCGACACAAGAAAATG<br>RV: GTGCAACTTTGGAGGATAGA | 300 nM |
| *CDKN1A* | FW: TGTCACTGTCTTGTACCCTTGT<br>RV: GCCGGCGTTTGGAGTGGTAG | 300 nM |
| *CDKN1B* | FW: ACTCTGAGGACACGCATTTGGT<br>RV: TCTGTTCTGTTGGCTCTTTTGTT | 300 nM |
| *MCL1* | FW: GTAATAACACCAGTACGGACGG<br>RV: TCCCGAAGGTACCGAGAGAT | 300 nM |
| *MP1LC3B* | FW: AAGGCGCTTACAGCTCAATG<br>RV: CTGGGAGGCATAGACCATGT | 150 nM |
| *PMAIP1* | FW: CGCGCAAGAACGCTCAACC<br>RV: CACACTCGACTTCCAGCTCTGCT | 300 nM |
| *ULK1* | FW: CAGACAGCCTGATGTGCAGT<br>RV: CAGGGTGGGGATGGAGAT | 150 nM |
| *ACTB* | FW: AGGCCAACCGCGAGAAG<br>RV: ACAGCCTGGATAGCAACGTACA | 150 nM |
| *HPRT1* | FW: GAACGTCTTGCTCGAGATGTGA<br>RV: TCCAGCAGGTCAGCAAAGAAT | 150 nM |

**Supplementary Table 5.** Cell cycle-, apoptosis-, and autophagy-related genes modulated by THZ-P1-2 in acute leukemia cells.

| Gene | MV4-11 cells | | | OCI-AML3 cells | | | Jurkat cells | | | NALM6 cells | | |
|---|---|---|---|---|---|---|---|---|---|---|---|---|
| | **FC**[1] | **S.D.** | ***p***[2] | **FC**[1] | **S.D.** | ***p***[2] | **FC**[1] | **S.D.** | ***p***[2] | **FC**[1] | **S.D.** | ***p***[2] |
| *CCNA2* | 0.57 | 0.08 | **0.002** | 0.78 | 0.04 | **0.0016** | 1.06 | 0.49 | 0.8178 | 1.33 | 0.15 | **0.021** |
| *CCNB1* | 0.89 | 0.19 | **0.0009** | 0.89 | 0.10 | 0.1082 | 0.97 | 0.19 | 0.7479 | 1.22 | 0.02 | **0.0003** |
| *CCND1* | 0.31 | 0.05 | **0.0001** | 0.91 | 0.09 | 0.1346 | 4.33 | 1.71 | **0.0298** | n.d. | n.d. | > 0.05 |
| *CCNE1* | 0.96 | 0.12 | 0.5821 | 0.95 | 0.19 | 0.6258 | 1.29 | 0.10 | **0.0117** | 1.19 | 0.02 | **0.0004** |
| *CDKN1A* | 0.98 | 0.13 | 0.7495 | 4.48 | 0.42 | **0.0005** | 4.47 | 1.88 | **0.0345** | 0.59 | 0.09 | **0.0029** |
| *CDKN1B* | 2.43 | 0.70 | **0.0268** | 2.39 | 0.11 | **0.0001** | 2.45 | 0.16 | **0.0004** | 2.23 | 0.25 | **0.0023** |
| *BCL2* | 1.10 | 0.07 | 0.0626 | 0.87 | 0.09 | 0.1848 | 1.73 | 0.24 | **0.0088** | 0.44 | 0.02 | **< 0.0001** |
| *BCL2L1* | 1.68 | 0.15 | **0.0028** | 1.75 | 0.04 | **< 0.0001** | 1.38 | 0.34 | 0.1066 | 1.91 | 0.14 | **0.0009** |
| *MCL1* | 1.70 | 0.30 | **0.0186** | 1.80 | 0.26 | **0.009** | 1.85 | 0.33 | **0.0141** | 1.40 | 0.22 | **0.0391** |
| *BAX* | 1.36 | 0.13 | **0.0113** | 2.42 | 0.24 | **0.0013** | 1.60 | 0.11 | **0.0017** | 1.38 | 0.18 | **0.0235** |
| *BAD* | 1.33 | 0.21 | 0.0546 | 1.07 | 0.19 | 0.5175 | 2.39 | 1.04 | 0.0749 | 1.16 | 0.31 | 0.3674 |
| *BCL2L11* | 2.92 | 0.22 | **0.0004** | 2.00 | 0.08 | **0.0001** | 2.04 | 0.22 | **0.0025** | 1.66 | 0.14 | **0.0026** |
| *BBC3* | 1.89 | 0.30 | **0.0098** | 5.20 | 0.44 | **0.0003** | 5.75 | 2.19 | **0.0227** | 2.07 | 0.39 | **0.0123** |
| *PMAIP1* | 0.53 | 0.22 | **0.0125** | 1.11 | 0.18 | 0.3208 | 1.50 | 0.72 | 0.2572 | 1.51 | 0.20 | **0.0146** |
| *GADD45A* | 1.24 | 0.23 | 0.1095 | 3.29 | 0.13 | **< 0.0001** | 2.85 | 0.30 | **0.0011** | 1.57 | 0.20 | **0.0109** |
| *ULK1* | 1.74 | 0.17 | **0.003** | 2.69 | 0.16 | **0.0002** | 1.10 | 0.08 | 0.084 | 1.70 | 0.18 | **0.0047** |
| *MAP1LC3B* | 2.54 | 0.42 | **0.0051** | 4.43 | 0.22 | **< 0.0001** | 2.23 | 0.25 | **0.0023** | 1.88 | 0.27 | **0.0075** |
| *BECN1* | 1.83 | 0.34 | **0.016** | 1.34 | 0.20 | **0.0449** | 0.98 | 0.19 | 0.8227 | 1.31 | 0.23 | 0.0773 |
| *BNIP3* | 1.41 | 0.28 | 0.0638 | 1.54 | 0.16 | **0.0067** | 1.42 | 0.13 | **0.0075** | 1.41 | 0.20 | **0.0258** |
| *ATG5* | 1.70 | 0.24 | **0.0108** | 2.09 | 0.23 | **0.0025** | 2.08 | 0.57 | **0.0317** | 1.54 | 0.20 | **0.013** |
| *ATG7* | 1.63 | 0.17 | **0.0055** | 2.99 | 0.32 | **0.0011** | 0.61 | 0.04 | **0.0003** | 0.97 | 0.05 | 0.2926 |
| *ATG10* | 1.17 | 0.12 | 0.0688 | 1.12 | 0.16 | 0.2218 | 0.56 | 0.10 | **0.0031** | 1.07 | 0.24 | 0.6034 |

Abbreviation: F.C., Fold-change; S.D., standard deviation; n.d., not detected.

[1]Fold-change of vehicle-treated cells.

[2]Student *t* test.
